# Contagious Antibiotic Resistance: Plasmid Transfer Among Bacterial Residents of the Zebrafish Gut

**DOI:** 10.1101/2020.11.09.375964

**Authors:** Wesley Loftie-Eaton, Angela Crabtree, David Perry, Jack Millstein, Barrie Robinson, Larry Forney, Eva Top

## Abstract

By characterizing the trajectories of antibiotic resistance gene transfer in bacterial communities such as the gut microbiome, we will better understand the factors that influence this spread of resistance. Our aim was to investigate the host network of a multi-drug resistance broad-host-range plasmid in the culturable gut microbiome of zebrafish. This was done through *in vitro* and *in vivo* conjugation experiments with *Escherichia coli* as donor of the plasmid pB10::*gfp*. When this donor was mixed with the extracted gut microbiome, only transconjugants of *Aeromonas veronii* were detected. In separate matings between the same donor and four prominent isolates from the gut microbiome, the plasmid transferred to two of these four isolates, *A. veronii and Plesiomonas shigelloides,* but not to *Shewanella putrefaciens* and *Vibrio mimicus*. When these *A. veronii and P. shigelloides* transconjugants were the donors in matings with the same four isolates, the plasmid now also transferred from *A. veronii* to *S. putrefaciens*. *P. shigelloides* was unable to donate the plasmid and *V. mimicus* was unable to acquire it. Finally, when the *E. coli* donor was added *in vivo* to zebrafish through their food, plasmid transfer was observed in the gut but only to *Achromobacter sp.,* a rare member of the gut microbiome. This work shows that the success of plasmid-mediated antibiotic resistance spread in a gut microbiome depends on the donor-recipient species combinations and therefore their spatial arrangement. It also suggests that rare gut microbiome members should not be ignored as potential reservoirs of multi-drug resistance plasmids from food.

**Importance:** To understand how antibiotic resistance plasmids end up in human pathogens it is crucial to learn how, where and when they are transferred and maintained in members of bacterial communities such as the gut microbiome. To gain insight into the network of plasmid-mediated antibiotic resistance sharing in the gut microbiome, we investigated the transferability and maintenance of a multi-drug resistance plasmid among the culturable bacteria of the zebrafish gut. We show that the success of plasmid-mediated antibiotic resistance spread in a gut microbiome can depend on which species are involved, as some are important nodes in the plasmid-host network and others dead-ends. Our findings also suggest that rare gut microbiome members should not be ignored as potential reservoirs of multi-drug resistance plasmids from food.

## 1. Introduction

Today many bacterial pathogens responsible for nosocomial infections are resistant to most if not all available antibiotics (Kåhrström 2013; World Health Organization. 2018). In contrast to the late 1960s when the US Surgeon General stated that it is “time to close the book” on infectious diseases, we are now warned by authorities such as the WHO and NIAID about an emanating ‘post-antibiotic’ era (World Health Organization 2014). What was underestimated early on was the ability of bacteria to rapidly adapt to selective pressures such as those created by the prolific use of antibiotics. One important mechanism of bacterial adaptation to antibiotics is the acquisition of antibiotic resistance genes from other, even distantly related bacteria through horizontal gene transfer (HGT) (Broaders *et al.*, 2013). One of the main HGT mechanisms responsible for antibiotic resistance spread to pathogens is conjugation (Carattoli, 2013; Mathers *et al.* 2015). Conjugation requires cell-to-cell contact and allows transfer of plasmid DNA from a donor to a recipient cell. To slow down the spread of antibiotic resistance we need to understand the bacterial reservoirs of the resistance genes, and how and where pathogenic bacteria acquire these genes from these reservoirs by conjugation (Andersson and Hughes, 2012; Piddock 2017).

One environment where antibiotic resistance genes are likely exchanged between resident and transient bacteria is the gastrointestinal tract of humans and animals. The gut is expected to be favorable to HGT for multiple reasons (Aminov, 2011). It provides near continuous nutrition to the gut microbiota and environmental conditions that allow for bacterial growth and high population densities that promote efficient conjugation. Plasmid transfer has been demonstrated in the mouse gastrointestinal tract (for example Licht *et al.*, 1999; García-Quintanilla *et al.*, 2008), in the guts of fleas and houseflies (Hinnebusch *et al.*, 2002; Fukuda *et al.*, 2015), and even in the gut of zebrafish (Fu *et al.*, 2017).

For quite some time there has been evidence for plasmid transfer in the human gut (Broaders *et al.*, 2013; Balis *et al.*, 1996; Datta *et al.*, 1981). Moreover, a comparison of 1,183 human associated bacteria and 1,052 bacteria from a broad range of environmental niches suggested that bacteria within the human gut microbiome may be 25-fold more likely to share genetic material than bacteria from other environments (Smillie *et al.*, 2011). Moreover, not only do plasmids carry accessory genes that encode antibiotic resistance but they also encode various pathogenicity factors (see Ogilvie *et al.*, 2012 for a comprehensive review) and genes that confer the ability to metabolize complex nutrients and degrade xenobiotic compounds. Due to their prevalence and potentially high rate of transfer in the gut, plasmids may provide functional redundancy to prevent the loss of key functions (Ogilvie *et al.*, 2012). However, such a functionally redundant network of mobile genetic elements could also lead to antibiotic resistance gene reservoirs that persist in the absence of antibiotic selection.

To understand how antibiotic resistance plasmids end up in human pathogens it is crucial to learn how, where and when they are transferred and maintained in members of bacterial communities such as the gut microbiome. One desirable model system for such studies are Zebrafish (*Danio rerio*) because they have a well-defined and comparatively simple core microbiome, and a digestive tract that is similar in organization and function to that of mammals. Therefore, they have been used to investigate host-microbe interactions, gut colonization and differentiation (Burns *et al.*, 2016; Lan and Love, 2012; Russo *et al.*, 2015; Stephens *et al.*, 2016), and recently also plasmid transfer (Fu *et al.*, 2017). To gain insight to the network of plasmid-mediated antibiotic resistance sharing in the gut microbiome, we investigated the transferability and maintenance of the multi-drug resistance (MDR) plasmid pB10::*gfp* among the culturable bacteria of the zebrafish gut microbiome, through both *in vitro* and *in vivo* studies. This plasmid belongs to the incompatibility group IncP-1, a well-known group of plasmids that are self-transmissible to a broad host range (BHR) of bacteria and likely to be involved in the spread of antibiotic resistance (Popowska 2013). In *in vitro* matings the efficiency of plasmid transfer was found to depend on the combination of bacterial species acting as plasmid donors and recipients. In our *in vivo* study only one particular species of the zebrafish gut microbiome effectively received and maintained the plasmid even though it constituted a small fraction of that community. Given conjugation requires cell contact, our findings suggest that the successful spread and persistence of a plasmid in a gut microbiome depends on the bacterial community composition and the spatial arrangement of its members. They also caution against using *in vitro* conjugation results to identify the likely reservoirs of MDR plasmids in a gut microbiome, as these can be rare community members.

## 2. Results

### 2.1 Plasmid transfer to culturable bacteria from the zebrafish gut microbiome

First we assessed the ability of plasmid pB10::*gfp* to transfer from an *E. coli* donor to bacteria of the zebrafish gut microbiome. We performed these conjugation experiments on an agar surface using *E. coli* (pB10::*gfp*) as a plasmid donor to either (i) the entire microbiome isolated from the zebrafish intestinal tract or (ii) four numerically dominant species that had been isolated from these gut microbiomes.

Conjugation experiments on R2A agar between *E. coli* AT1036 (pB10::*gfp*) and the zebrafish gut microbiome (approximately 1 × 10^8^ culturable bacteria) yielded pB10::*gfp*-containing microbiome members (called transconjugants) at a frequency of 1.5 (±1.5) × 10^−4^ per donor. For simplicity we often refer to this transconjugant/donor ratio from here on as the ‘transfer frequency’. Based on analysis of their 16S rRNA gene sequences these transconjugants all belonged to a single species, *Aeromonas veronii*. For the second *in vitro* method we first isolated individual bacterial strains from the combined guts of four zebrafish, using different growth media. Based on differences in colony morphology we isolated and purified twenty nine strains. Based on their 16S rRNA gene sequences, they belonged to the four gamma-Proteobacterial species *Aeromonas veronii, Plesiomonas shigelloides, Shewanella putrefaciens* and *Vibrio mimicus*. Conjugation experiments done using the *E. coli* (pB10::*gfp*) donor and these four species as recipients yielded transconjugants for *A. veronii* and *P. shigelloides* at frequencies of 8.8 (±5.5) × 10^−4^ and (±0.1) × 10^−3^ per donor, respectively. In contrast, transfer to *V. mimicus* and *S. putrefaciens* could not be detected (<1 × 10^−8^ transconjugants per donor). The plasmid was thus able to transfer from an *E. coli* donor to at least two of the four dominant culturable members of the zebrafish gut microbiome.

### 2.2 Novel hosts may act as plasmid donor

Since two culturable zebrafish gut microbiome members received the antibiotic resistance plasmid from *E. coli,* we determined if they could further spread it to other species and whether various donor/recipient combinations would affect the efficiency of plasmid transfer. To do this, the *A. veronii* and *P. shigelloides* transconjugants were each employed as donors in conjugation assays with five recipients: *E. coli* EC100 and rifampicin resistant (RifR) mutants of each of the four gut isolates, *A. veronii*, *P. shigelloides, S. putrefaciens*, and *V. mimicus.* As shown in Table 1 the frequency of plasmid transfer to one particular recipient clearly depended on the identity of the donor, and the efficiency by which a donor transferred the plasmid depended on the recipient. Strikingly, though *P. shigelloides* was a good recipient, it was a bad plasmid donor as it was unable to transfer the plasmid to any of the five recipients. The lack of plasmid transfer from this host was not due to plasmid integration in the chromosome as extrachromosomal plasmid DNA was visualized on an agarose gel (Fig. 2). What is also clear from Table 1 is that *A. veronii* but not *E. coli* was able to transfer the plasmid to *S. putrefaciens*, albeit still at a low frequency, showing a clear donor effect. We were unable to use *S. putrefaciens* as a donor in these reciprocal transfer experiments since it was intrinsically resistant to both rifampicin (Rif) and nalidixic acid (Nal), which precluded distinguishing between donors and transconjugants. Transfer to *V. mimicus* could not be detected at all, making it also impossible to use it as a plasmid donor. These results suggest that the trajectory of spread of a BHR MDR plasmid in a microbiome is determined by the identity of both the donor and recipient.

**Fig. 1.**
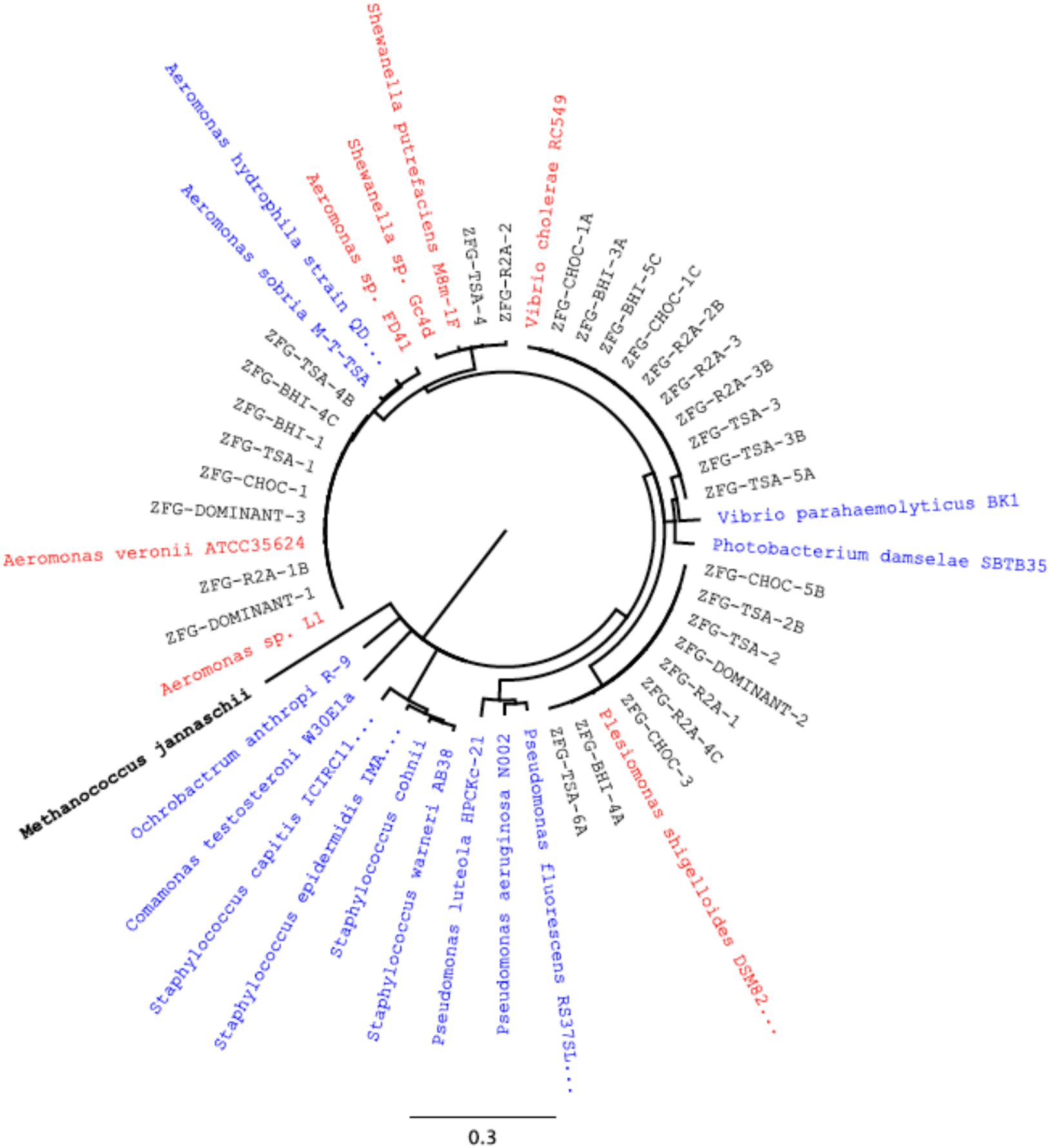
All 29 unique zebrafish gut isolates grouped into four different genera based on their 16S rRNA gene sequences: *Aeromonas, Plesiomonas, Vibrio* and *Shewanella*. ZFG, bacterial strains isolated from the zebrafish gut; DOMINANT, numerically dominant colony morphology types, originally identified on R2A agar, and which successfully acquired the plasmid following conjugation with the donor *E. coli*; BHI, CHOC, R2A and TSA, abbreviations of media used to isolate the strains (see Materials and Methods); Red, culture collection strains most similar to isolates; Blue, zebrafish core microbiome reference strains (Cantas *et al.*, 2012); The tree was rooted using the Archaean *Methanococcus jannaschii*.

**Fig. 2.**
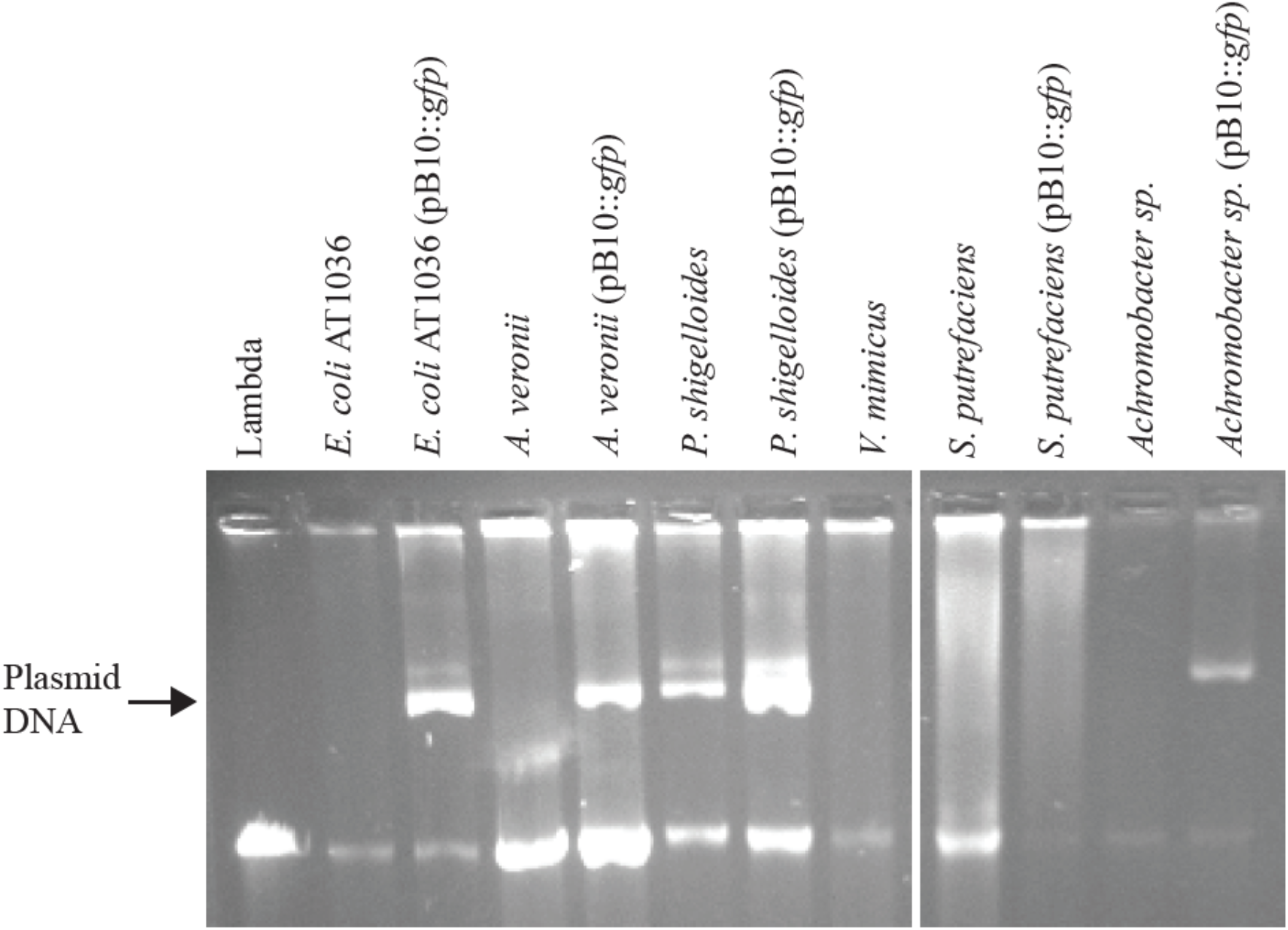
Agarose gel electrophoresis of plasmid DNA extracted from plasmid-free and plasmid-containing strains.

**Table 1.** Plasmid transfer frequencies^*^ in reciprocal matings demonstrate the importance of both donor and recipient identity in the efficiency of plasmid transfer.

|  | Donor of pB10::gfp |  |  |  |
| --- | --- | --- | --- | --- |
|  |  | *E. coli* AT1036 | *A. veronii* | *P. shigelloides* |
| **Recipient** | *E. coli* EC100 Nal<sup>R</sup> | $2.8 (\pm 1.4) \times 10^{-2}$ | $8.8 (\pm 5.5) \times 10^{-4}$ | ND |
| | *A. veronii* Rif<sup>R</sup> | $3.4 (\pm 0.7) \times 10^{-3}$ | $2.9 (\pm 0.2) \times 10^{-1}$ | ND |
| | *P. shigelloides* Rif<sup>R</sup> | $1.5 (\pm 0.0) \times 10^{-4}$ | $2.2 (\pm 1.0) \times 10^{-3}$ | ND |
|  | *V. mimicus* Rif<sup>R</sup> | ND** | ND | ND |
| | *S. putrefaciens* | ND | $4.2 (\pm 3.2) \times 10^{-5}$ | ND |

\* Frequencies are expressed as transconjugants per donor after the matings (n = 3).
\*\* ND, transconjugants were below the detection limit of $10^{-8}$ .

### 2.3 *The persistence of pB10::*gfp *in zebrafish gut bacteria was host-dependent*

For a microbiome member to be an important reservoir of an MDR plasmid, it must not only acquire an incoming plasmid but also retain it sufficiently long in the absence of selection for plasmid-encoded genes. To determine whether pB10::*gfp* was able to persist in our *A. veronii*, *P. shigelloides* and *S. putrefaciens* isolates, we monitored the fraction of plasmid-containing cells in serially transferred populations in the absence of antibiotic selection for 10 days (100 generations). The plasmid was very persistent in *P. shigelloides* but much less so in the other two strains (Fig. 3). Thus, although *A. veronii* was both a rather good recipient and donor of the plasmid (Table 1), it was not good at maintaining the plasmid and may thus represent only a transient host. In contrast *P. shigelloides* was very good at retaining the plasmid but unable to transfer it to other bacteria, suggesting a dead-end for the plasmid in its transmission network. Finally, *S. putrefaciens* was very poor both at receiving and retaining the plasmid. These results show that persistence of the plasmid was variable in zebrafish gut bacteria and did not necessarily correlate with their host’s ability to receive or further transfer the plasmid.

**Fig. 3.**
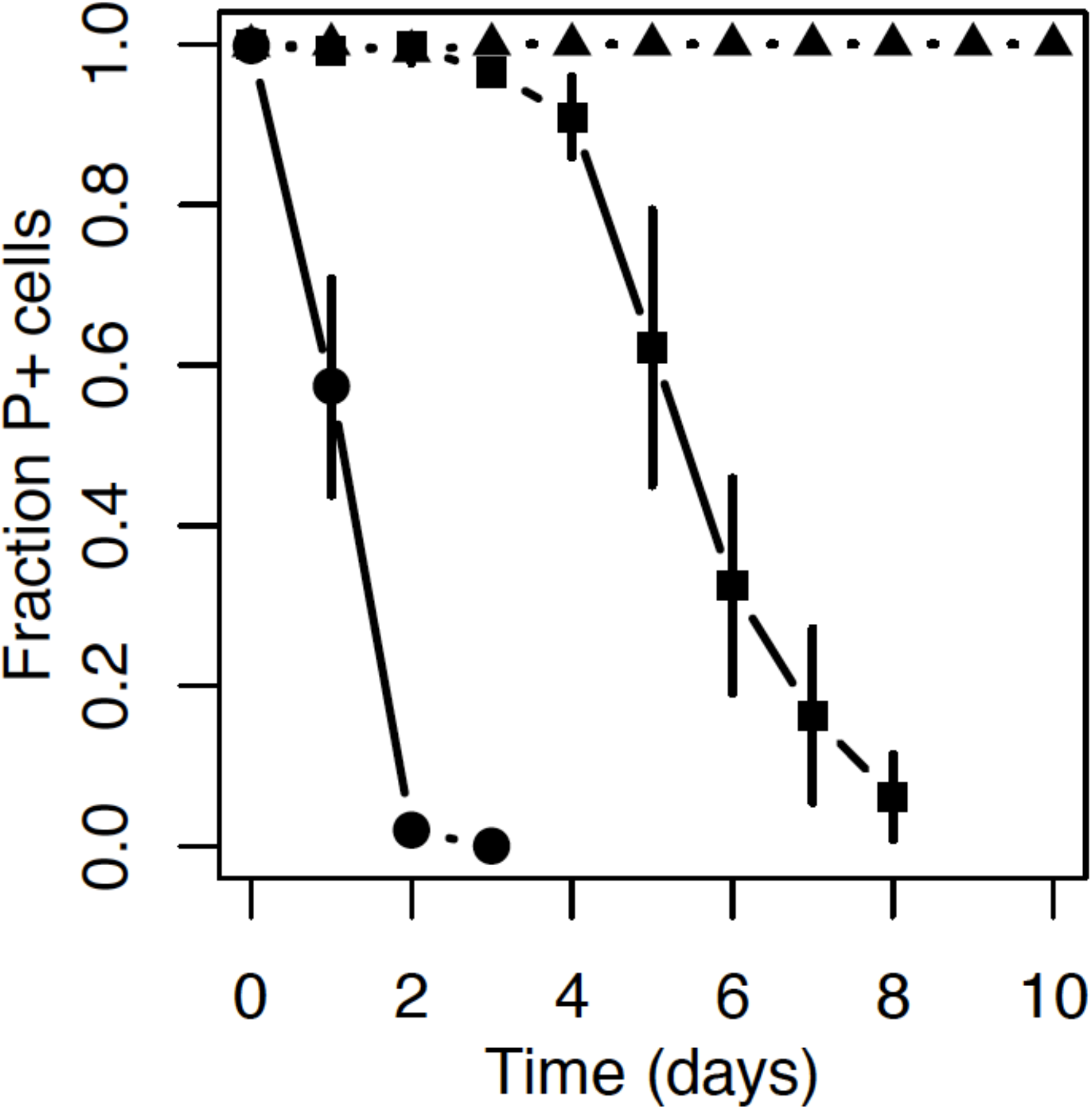
The persistence of plasmid pB10:: *gfp* in zebrafish gut isolates over 100 generations of growth in the absence of antibiotic selection varied greatly from high persistence (*P. shigelloides;* triangles) to moderate (*A. veronii*; squares) and poor persistence (*S. putrefaciens*; circles).

### 2.4 *Plasmid transfer* in vivo

Next we examined if the plasmid could transfer to bacteria in the gut of zebrafish exposed to tetracycline, one of the antibiotics for which the plasmid encodes resistance and which is frequently used in aquaculture (Tuševljak *et al.*, 2013). Briefly, with 32 fish divided over eight separate tanks, half of the fish were fed twice daily with food pre-mixed with the plasmid donor (treated; tanks A to D), while the other half received food pre-mixed with an isogenic plasmid-free strain (untreated; tanks E to H). After analyzing the gut microbiomes of the eight fish populations harvested on the 23^rd^ day, 23 green fluorescent transconjugants were observed on transconjugant-selective agar plates at an average frequency of 1.2 (± 0.8) per 10^6^ culturable gut bacteria from the zebrafish in three of the four treated tanks. In contrast, no green fluorescent strains were detected in the guts of the untreated groups. No fluorescent *E. coli* AT1036 (pB10::*gfp*) donors were detected on donor-permissive agar plates, verifying that this *E. coli* strain was incapable of establishing itself in the zebrafish gut. All 23 transconjugant colonies looked identical and, based on comparisons of a 1.3-kb fragment of their 16S rRNA gene sequences, they all belonged to the genus *Achromobacter* (Fig. 4). Since plasmid donors were no longer fed to the fish during the last two days before harvesting the microbiomes, these transconjugants were present in the zebrafish gut for at least two days post treatment.

**Fig. 4.**
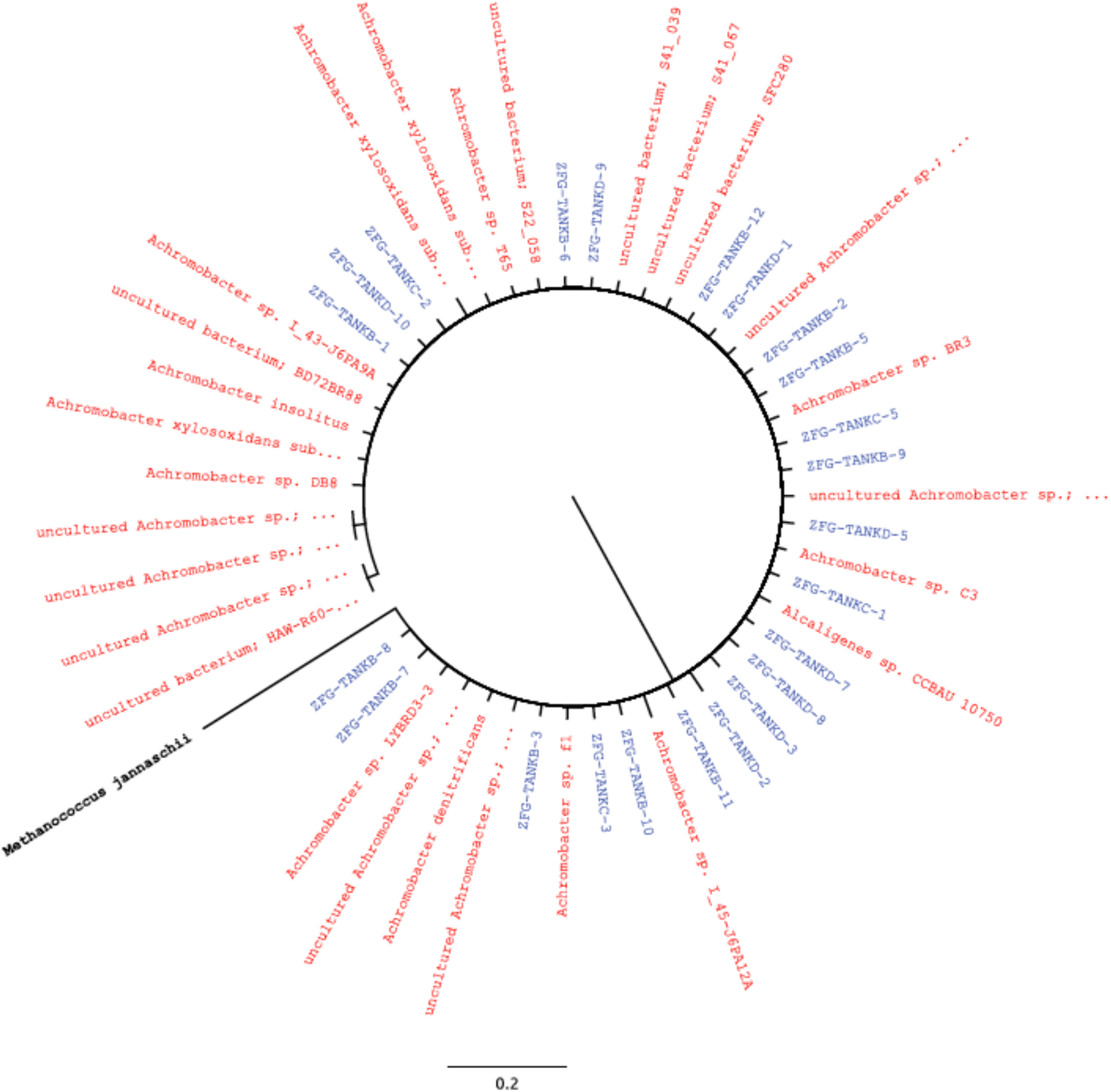
Phylogenetic diversity of 23 *Achromobacter sp*. transconjugants isolated from the guts of zebrafish (ZFG, blue) (3 of 4 tanks) after 21 days of repeated inoculation with the plasmid donor. Strains most similar to these isolates are shown in red. The tree was rooted using the Archaean *Methanococcus jannaschii*.

Since *Achromobacter* is generally described as a common water-borne organism (Garrity *et al.*, 2005) and has not previously been referenced as a resident of the zebrafish gut microbiome, we determined if *Achromobacter sp.* was indeed present within the guts of our zebrafish populations. This was done by constructing and sequencing 16S rRNA amplicon libraries using gDNA isolated from the gut microbiome samples. The DNA sequence analysis showed that *Achromobacter sp.* was present at a low frequency in the gut microbiome of both the treated and untreated populations [7.9 (± 4.2) × 10^−3^% and 8.2 (± 5.6) × 10^−2^%, respectively] (Fig.5). In comparison, *Aeromonas* represented 40.3 ± 27.1 % and 16.2 ± 5.0% of the gut microbiomes of treated and untreated populations, respectively. Despite being the most numerically dominant genus in these guts and a good plasmid recipient *in vitro* (section 2.1), no transconjugants of this genus were identified at the end of this *in vivo* experiment. Our findings suggest that factors other than species abundance determine the *in vivo* trajectories of plasmid transfer and establishment in a gut microbiome.

**Fig. 5.**
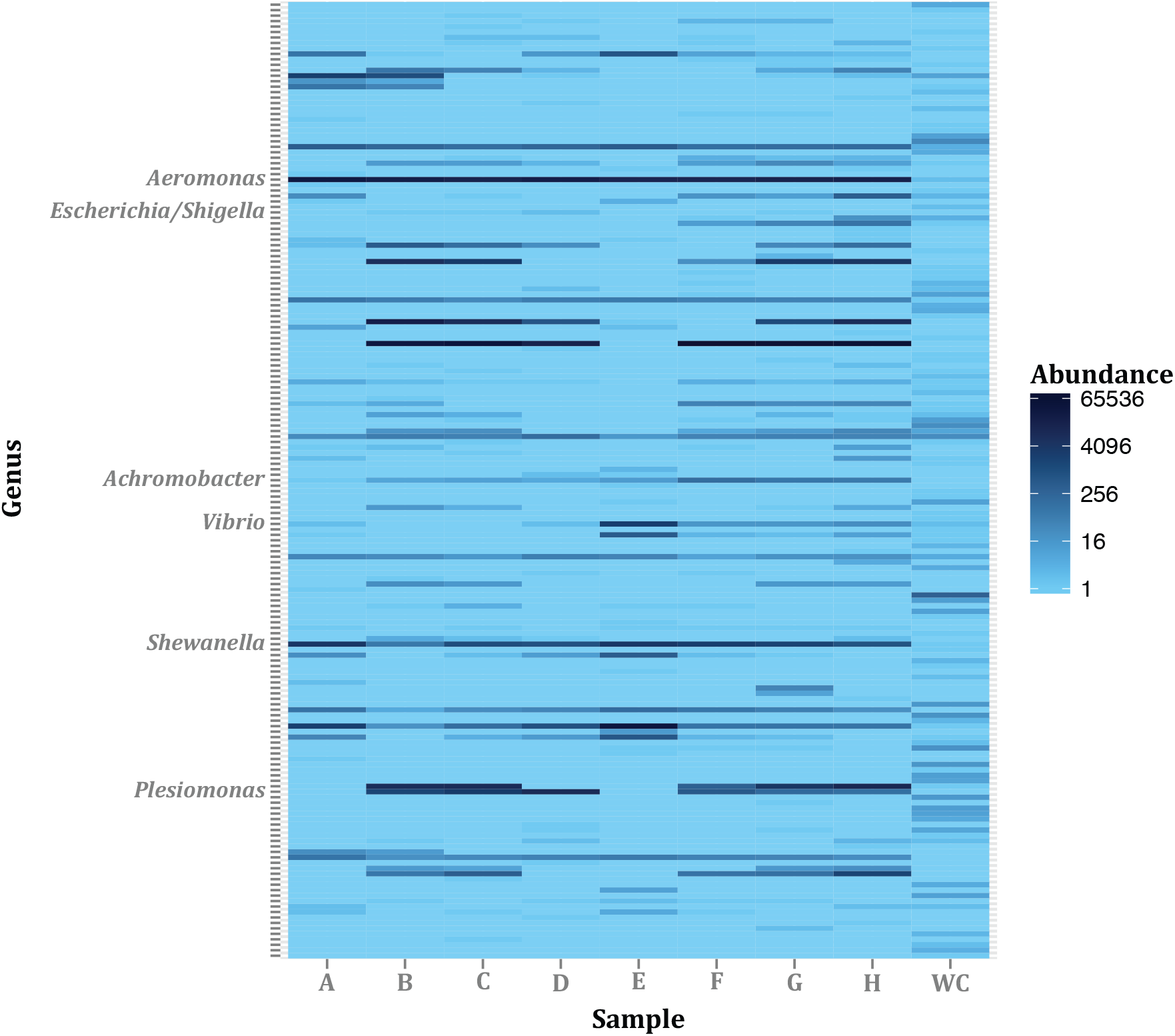
*Achromobacter sp*. was present at low abundance among the many genera in the gut microbiota of the treated zebrafish populations (A to D: 7.9 (± 4.2) × 10^−3^ %) and untreated populations (E to H: 8.2 (± 5.6) × 10^−2^ %), and not detectable in the water control (WC). Only the relevant cultured genera or operational taxonomic units (OTUs) are indicated. Abundance is expressed in 16S rRNA gene sequence reads. A complete list of all the OTUs can be found in Table S1.

### 2.5 *Transferability to and persistence of pB10::*gfp *in* Achromobacter sp

To determine if a high frequency of transfer from *E. coli* to *Achromobacter sp.* could in part explain why *Achromobacter sp.* was the only species of *in vivo* transconjugants detected, we determined the transfer frequency *in vitro*. On an agar surface, plasmid pB10::*gfp* transferred from *E. coli* AT1036 to *Achromobacter sp.* at a frequency of 4.3 (± 2.0) × 10^−2^ transconjugants.donor^−1^. When comparing this to plasmid transfer under the same conditions from the same *E. coli* to other species used in this study (Table 1), this frequency was on average 13 times higher compared to transfer to *A. veronii*, 287 times higher than to *P. shigelloides*, and similar to the frequency of transfer between two isogenic *E. coli* strains. Thus, even though *Achromobacter sp.* was present at only 0.008 % of the gut microbiome, its high proficiency as a recipient for pB10::*gfp* likely allowed it to acquire that plasmid within the zebrafish gut.

For gut bacteria to become new reservoirs of horizontally acquired MDR plasmids, they not only need to receive the plasmid, but also retain it under conditions of low or no antibiotic pressure. Therefore, we also measured the persistence of pB10::*gfp* in *Achromobacter sp.* in serial batch culture. After approximately 100 generations of growth in the absence of antibiotic selection for plasmid maintenance, about 80% of the *Achromobacter sp.* population still retained the plasmid (Fig. 6). Since some bacterial strains have shown no loss of plasmid pB10::*gfp* in this time frame and others a much more rapid loss (De Gelder *et al.*, 2007), this *Achromobacter sp.* strain seems to be a moderately good host for this plasmid. A combination of a high plasmid transfer frequency and moderate plasmid persistence likely explained how this plasmid-host pair formed in the gut and then persisted for at least two days since the last *E. coli* donor cells were added.

**Fig. 6.**
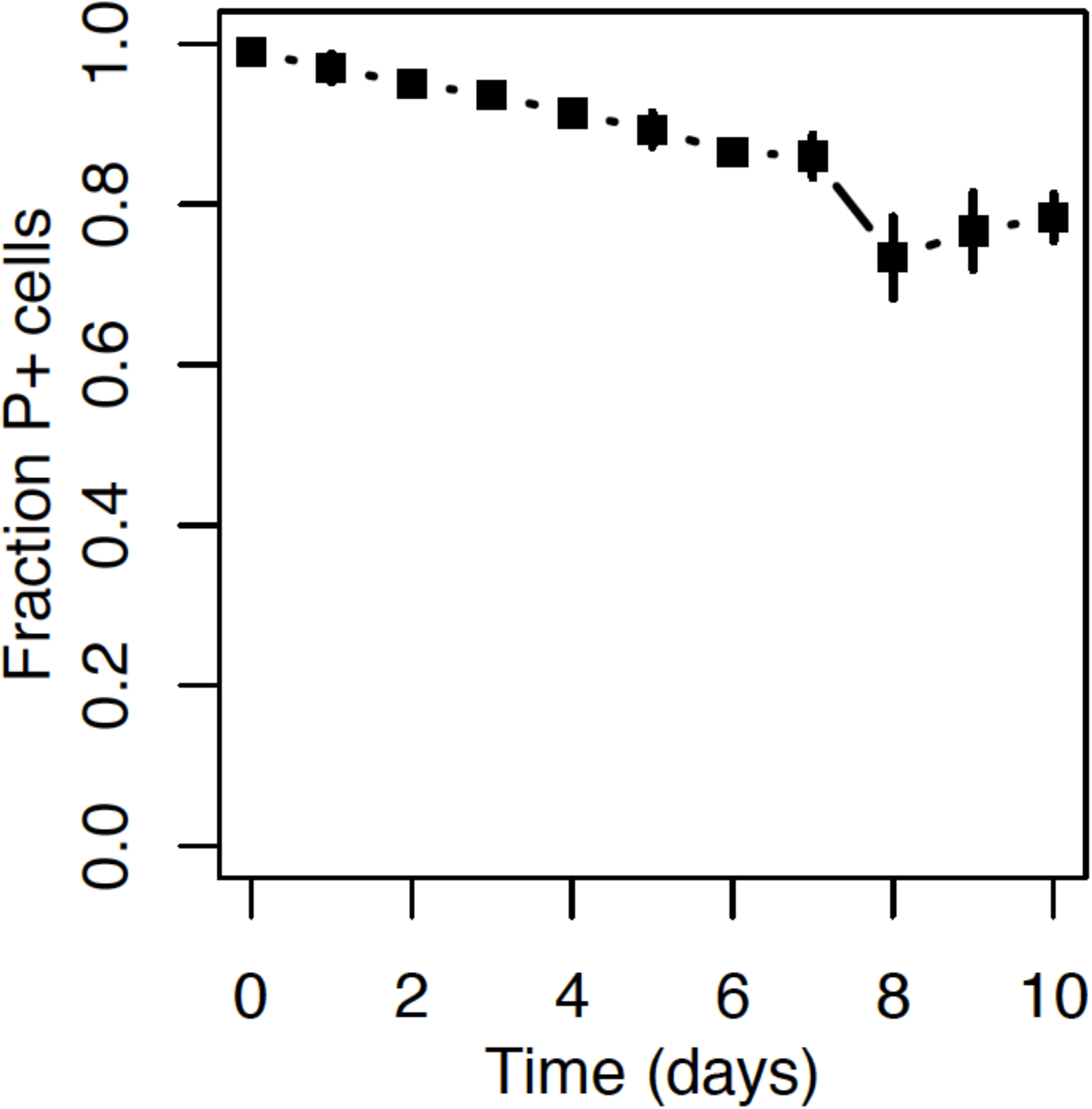
Plasmid pB10::*gfp* is moderately persistent in *Achromobacter sp*.

## 3. Discussion

If we want to slow down the alarmingly rapid spread of resistance to critically important antibiotics we need to better understand the plasmid transfer networks that facilitate this spread. Conjugation of self-transmissible plasmids is likely a major pathway for horizontal gene transfer among bacteria, and so-called ‘epidemic’ plasmids play a critical role in global resistance spread, in particular among multidrug-resistant *Enterobacteriaceae* (Carattoli, 2013; Mathers *et al.*, 2015). Using zebrafish as a model system we showed here that an MDR plasmid introduced with the fish food can transfer and establish itself in one of the quite rare gut microbiome members, creating a new reservoir of mobile resistance genes. We also showed that key factors to this successful spread are likely the efficiency of plasmid transfer, which itself depends on the combination of donor and recipient bacteria in the transfer network, and the strains’ ability to maintain an MDR plasmid in the absence of antibiotics.

The gut microbiome of zebrafish appears to be relatively conserved, with a small group of core genera that are present in most sampled zebrafish (Roeselers *et al.*, 2011), including those in this study (Table S1). Of the most abundant bacterial genera in our zebrafish microbiomes, *Aeromonas, Shewanella, Vibrio*, and *Plesiomonas*, the first three are known to be part of this core microbiome (Roeselers *et al.*, 2011). Furthermore, most of the bacteria present in the gut belonged to the Gram-negative phylum Proteobacteria. This group of bacteria actively participates in HGT (Kloesges *et al.*, 2011) and is within the host range of several MDR plasmid families including the IncP-1 plasmids such as our model plasmid pB10 (Suzuki *et al.*, 2010). The phylum Proteobacteria also contain several human pathogens listed by the WHO in 2017 as being of priority 1 (‘critical’) (World Health Organization 2017). The gut microbiome of zebrafish is thus an ideal model system for research pertaining to the transfer and maintenance of MDR plasmids in microbial communities.

We clearly showed that the success of plasmid transfer between bacterial isolates from the gut microbiome depends on the identity of both the donor and recipient. Some hosts were good recipients but poor donors or *vice versa*, while others were both good donors and recipients. This is in line with a previous study from our group showing that the donor species defined the host range of pB10 within a wastewater activated sludge community (De Gelder *et al.*, 2005). It is also consistent with an *in vitro* conjugation study by Dionisio *et al.* (2002), who showed a significant difference among enterobacterial species and even *E. coli* strains in the ability to donate the F-type plasmid R1. They postulated that the best donor strains can act as ‘amplifiers’ in a community and thereby facilitate the spread of antibiotic resistance We confirm here that some strains can be important nodes in the plasmid transfer network while others may be dead-ends. To slow down the spread of antibiotic resistance it is important to identify these critical nodes in microbiomes as well as the molecular mechanisms underlying their proficiency as plasmid donor or recipient.

One striking finding of our study was the inability of *P. shigelloides* to transfer the plasmid to any of the five different species tested, in spite of it being a rather good recipient when combined with each donor. This seems to be a clear example of a dead-end species in the transfer network. This poor conjugation proficiency could be due to inhibition of conjugation by another resident plasmid. Fig. 2 shows that it was the only host showing a second plasmid DNA band on agarose gel after plasmid extraction. Negative regulatory effects of co-resident plasmids on the conjugative transfer of a specific plasmid have been long known (Datta *et al.*, 1971). It is referred to as fertility inhibition and has previously been demonstrated for IncP-1 plasmids like pB10 (Santini and Stanisish, 1998). Dioinisio *et al.* (2002) also provided some evidence that plasmid-encoded fertility inhibition systems like FinOP on plasmid R1 were involved in the variable ability to serve as a donor, likely due to interaction with native plasmids of these strains. This phenomenon has also been more recently described for several plasmid combinations (Gama *et al.*, 2017), but for many the molecular mechanisms remain elusive. Whatever the mechanism here, given that the combination of donor and recipient hosts determines successful plasmid transfer, the transferability of a plasmid within a gut microbiome likely depends on who neighbors whom. Our findings suggest the gene transfer network might not only be determined by the composition of a bacterial community but also by the spatial arrangement of its members.

Transfer of a plasmid to a given host does not necessarily translate to successful establishment in that host (Bingle and Thomas, 2001; Adamczyk and Jagura-Burdzy, 2003). If the replication or partitioning systems do not function optimally, or if the cost of the plasmid is high the plasmid will fail to persist in that population (Ponciano *et al.*, 2007). This was shown here by the inability of the plasmid to persist in *A. veronii*. In contrast, the plasmid was highly persistent in *P. shigelloides* and persisted moderately well in *Achromobacter sp.* Thus, even in the absence of antibiotics, gastrointestinal bacteria that efficiently acquire and retain an MDR plasmid could ensure the persistence of that plasmid and its resistance genes within the gut microbiome. Importantly, the efficiency by which a host receives a plasmid does not necessarily correlate with its ability to subsequently retain it. This serves as a reminder that plasmid transfer from a donor, establishment in a recipient and its subsequent persistence are distinctly unique processes that all contribute to the success of plasmids in any microbiome.

The zebrafish gut microbiome members that were detected as new hosts of our BHR MDR plasmid differed between the *in vitro* and *in vivo* conjugation experiments. In *in vitro* matings between *E. coli* and the extracted zebrafish gut microbiome, the only species of transconjugants detected (*A. veronii*) was numerically dominant in this microbiome, both based on 16S rRNA gene amplicon sequencing (Fig. 5) and plate counts (Fig. 1). In matings between pure cultures it acquired the plasmid at a moderately high frequency compared to other tested species (Table 1). Consistent with this, Fu *et al.*, (2017) recently demonstrated that *Aeromonas* species represented the dominant fraction of the zebrafish gut microbiome based on a cultivation-independent methods. They also showed that members of this genus were common among the transconjugants of the IncP-1α plasmid they introduced in *in vivo* experiments. In contrast to the findings of Fu *et al.* (2017) and of our *in vitro* experiment, the only transconjugant we detected in our *in vivo* experiment was *Achromobacter sp.,* a minority member of the zebrafish gut microbiome (Fig. 5). Its low proportion in the gut community explains why it was not detected on any of the agar media, nor as a transconjugant in the *in vitro* mating with extracted gut microbiome, where the roughly 5,000-fold more dominant *A. veronii* apparently crowded the transconjugant plates. *Achromobacter* was also present in the gut of the zebrafish used by Fu *et al.*, (2017), varying in abundance along the fore-, mid- and hindgut, but it was not shown to acquire their resistance plasmid. There could be several possible reasons for this discrepancy, none of which are mutually exclusive: i) differences in plasmid donor strains and the way they were administered to the fish; ii) *Achromobacter* may not be a favorable host for IncP-1α plasmids used by Fu *et al.* as there is no complete overlap between the host ranges of IncP-1α and −βplasmids (pB10 in our study) (Norberg *et al.*, 2011), iii) the *Achromobacter.* strains in these two studies may have been distinctly different and plasmid host range is known to vary greatly between and even within species (De Gelder *et al.*, 2007), and iv) our cultivation-dependent technique had a better detection limit, allowing to identify transconjugants present at low abundance.

*Achromobacter sp.* was the best recipient in *in vitro* conjugation experiments between *E. coli* (pB10) and pure cultures of recipients. The plasmid was also rather persistent in this host, more so than in *A. veronii*. High plasmid transfer frequencies and plasmid persistence can satisfy the ‘existence conditions’ for plasmids in bacterial populations, as any plasmid loss can possibly be overcome by reinfection of the plasmid from neighboring cells (Stewart and Levin, 1977, Ponciano *et al.*, 2007). Using *in vitro* conjugation experiments to identify which microbiome members play an important role in the spread of a particular plasmid may thus be misleading, as these results may be determined by relative abundance and plasmid transfer frequency but not by plasmid persistence and *in vivo* conditions. Our findings uniquely emphasize that even rather rare species in a gut microbiome may become important reservoirs of antibiotic resistance if their ability to acquire resistance plasmids from bacteria introduced with food is high.

The detection of only one species in the zebrafish gut that received our plasmid in our in *vivo* experiments does not by any means imply that it was the only member that received the plasmid. It has been demonstrated that the host range of bacteria to which a plasmid can transfer exceeds the range in which they can replicate (Musovic *et al.*, 2006, Waters, 1999). Therefore some members like *A. veronii* here may have received the plasmid in the zebrafish gut, passed it on to others, and subsequently lost it. Other members were likely not culturable (Cantas *et al.*, 2012; Roeselers *et al.*, 2011, Fu *et al.*, 2017). Thus, the frequency and range of plasmid transfer is likely underestimated. We can also not exclude that conjugative transfer of the plasmid from the *E. coli* donor to *Achromobacter sp.* took place in the water environment prior to *Achromobacter* establishing in the gut. Irrespective of the route *Achromobacter sp.* transconjugants were present in the zebrafish gut for at least two days post donor treatment and are thus likely an important link in this plasmid transfer network.

While a full understanding of the plasmid transfer network in a gut microbiome will require cultivation-independent monitoring of introduced and indigenous plasmids with methods such as proximity ligation (Hi-C) (Marbouti *et al.* 2017; Stalder *et al.*, 2019), several important messages can be drawn from this cultivation-dependent study: i) successful transfer of the plasmid is very much dependent on the combination of donor and recipient, and therefore an MDR plasmid network in a non-well mixed system like the gut is in part defined by both the composition and spatial organization of the microbiome; ii) caution should be taken when drawing conclusions about the range of MDR plasmid spread from *in vitro* data, as *Achromobacter sp.* would have been missed here without the *in vivo* experiment, and iii) in spite of their low relative abundance, rare gut microbiome members could be important reservoirs of MDR plasmids introduced through food.

## 4. Materials and Methods

### 4.1 Bacterial strains, plasmids, media, and growth conditions

Our model plasmid was the BHR MDR plasmid of the IncP-1β family, pB10::*gfp*. This plasmid was previously constructed by inserting mini-Tn*5*-PA1– 04/03::*gfp*, a Tn*5* derivative transposon encoding green fluorescent protein (GFP), in the 68.3-kb plasmid pB10 (Van Meervenne *et al.* 2012). This plasmid is self-transmissible and codes for resistance to tetracycline, amoxicillin, sulfonamide, streptomycin, ionic mercury, and kanamycin (the latter encoded on the mini-Tn*5*). Antibiotics kanamycin (50 μg.ml^−1^) and tetracycline (10 μg.ml^−1^) were used to select for the plasmid.

Table 2 specifies relevant characteristics of each bacterial strain used in the study. *E. coli* AT1036 was cultured on Luria-Bertani (LB) or tryptic soy (TS) media at 30 °C, and diaminopimelic acid (DAP) was added to a final concentration of 100 μg.ml^−1^ when required. *Achromomacter sp., Aeromonas veronii, Plesiomonas shigelloides, Shewanella putrefaciencs and Vibrio mimicus* were cultured in TS at 26 °C. To obtain Rif resistant (RifR) mutants of the latter four strains they were serially sub-cultured in TS supplemented with increasing concentrations of rifampicin, from 20 to 200 μg.ml^−1^.

**Table 2.**
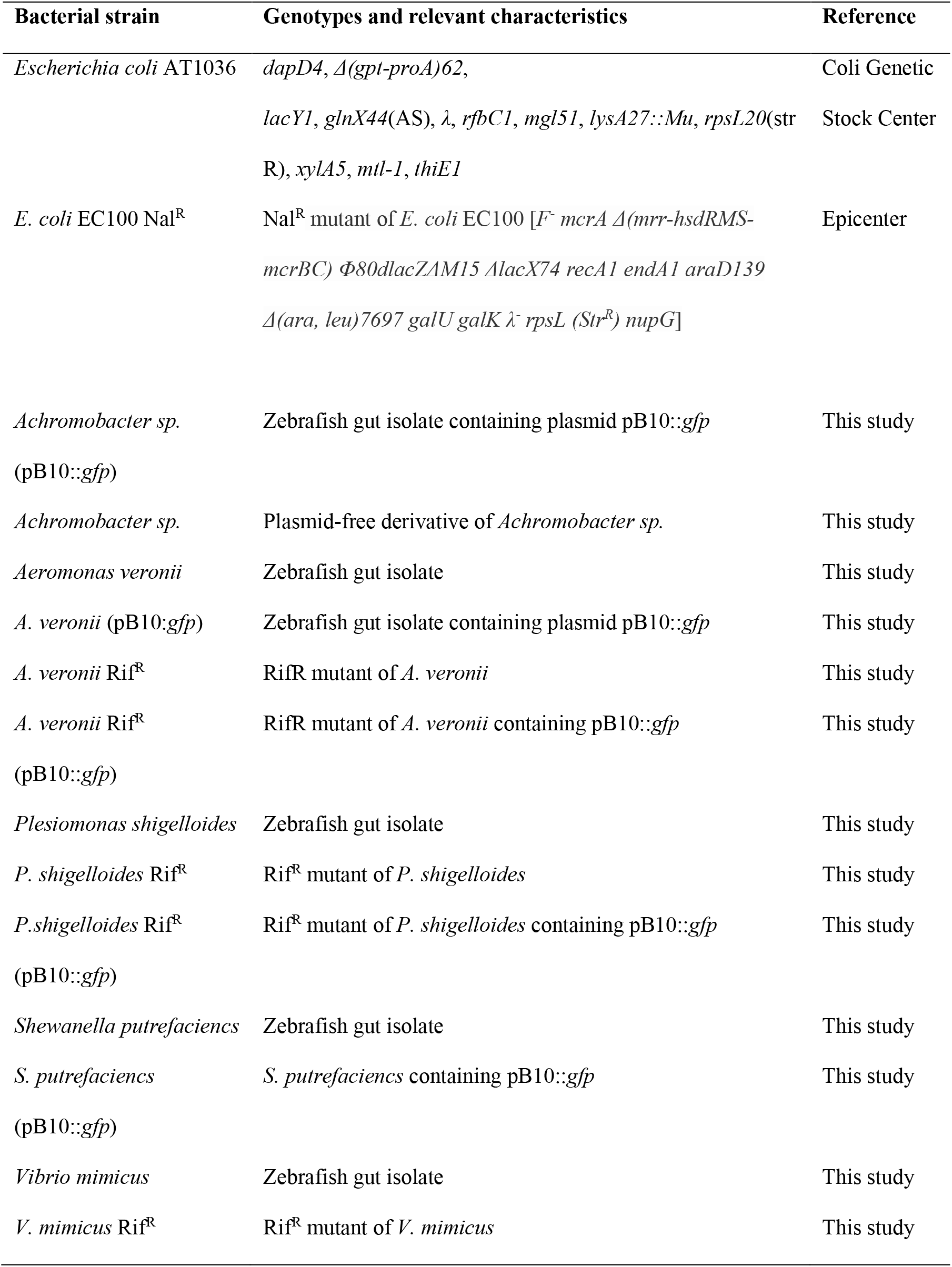
Bacterial strains used in this study.

### 4.2 General DNA manipulation techniques

Plasmid DNA was isolated and purified using a PureYield Plasmid Miniprep System (Promega) and gel electrophoresis were carried out using standard techniques (Sambrook and Russell, 2001). Genomic DNA (gDNA) was isolated using a Genelute bacterial gDNA kit (Sigma-Aldrich). All polymerase chain reaction (PCR) experiments, except those described in section 4.8, were performed using *Taq* Master Mix (NEB) as per the manufacturer’s instructions. The reaction parameters included an initial denaturation step of 10 min at 94°C, followed by 30 cycles of denaturation (30 s at 94°C), a variable annealing step dependent upon the average primer annealing temperature, and an elongation step at 72°C with the extension time depending on the amplicon size.

### 4.3 General zebrafish husbandry and gut bacteria isolation

Zebrafish (Scientific Hatcheries Inc) of unknown genetic background were maintained in a recirculating system (Aquaneering) at 28.5°C and fed twice daily with soy protein concentrate-based pellet food unless otherwise stated. Procedures involving animals were approved by the University of Idaho Institutional Animal Care and Use Committee. Prior to harvesting the gut bacteria, the fish were starved for 24 hours and individually anesthetized with 170 mg/l MS222 (Tricaine methanosulfonate; pH ∼7.0; Argent Laboratories). Each fish was aseptically dissected in a petri dish using a surgical blade. The entire for-, mid- and hind gut were placed into sterile 2 ml microfuge tubes and placed on ice. Disposable plastic inoculation loops were used to grind the guts and squeeze out the bacteria against the round bottom of the microfuge tube. To suspend the bacteria 2 ml of PBS (pH 7.4; 4 °C) was added and the samples were vortexed vigorously for 1 min. To separate the bacterial suspension and the gut lining, the samples were centrifuged at 800 rpm for 1 min and the bacteria-containing supernatant transferred to a sterile 2 ml microfuge tube. To collect the bacterial cells, the bacterial suspensions were centrifuged at 8000 rpm for 4 min and the pellet suspended in 2 ml PBS (pH 7.4; 4 °C). This procedure was repeated twice to wash the bacterial cells.

### 4.4 16S rRNA identity of clonal isolates

Individual bacterial colonies from the combined guts of four zebrafish were obtained by spreading 100 μl of a 10-fold dilution series onto Brain-Heart-Infusion (BHI), Reasoner’s 2A (R2A), TS and Chocolate agar and incubated at 26 °C for 48 hours. All media contained cycloheximide (100 μg.ml-1) to inhibit fungal growth. To obtain clonal isolates, a total of 29 colonies with unique morphologies were streaked onto their respective isolation media and TS agar (TSA). Single colonies were inoculated into 5 ml TS broth and grown overnight at 26 °C, and 1 ml of each culture was used for gDNA purification. 1 ng of gDNA was used as template for 16S rRNA gene amplification using the 27f (5’-AGAGTTTGATCMTGGCTCAG-3’) and 1492r (5’-TACGGYTACCTTGTTACGACTT-3’) 16S rRNA primers described by Roeselers *et al.* (2011). The ~1.6-kb PCR products were sequenced by Sanger sequencing at Elim Biopharmaceuticals Inc (California). The species identity of clonal isolates was determined by comparisons of their 16S rRNA gene sequences to known sequences within the RDP database. Phylogenetic trees were constructed using the default Alignment and Tree Builder functions within the Geneious R10 software package.

### 4.5 In vitro plasmid transfer

The original plasmid donor in the quantitative conjugation assays with zebrafish gut isolates as recipients was *E. coli* AT1036 (pB10::*gfp*), a DAP auxotroph. After these conjugation assays the resulting transconjugants served as donors in subsequent matings with Rif^R^ mutants of these isolates, and with Nal^R^ *E. coli* EC100.

To investigate plasmid transfer from the *E. coli* donor to the zebrafish gut microbial community *in vitro*, the recipient community was prepared in 1 ml PBS from the combined guts of four zebrafish as described in Section 4.3, and the conjugation assay was carried out on R2A. For all other *in vitro* conjugation experiments, the donors and recipients were prepared from cultures that were grown overnight at 30°C for *E. coli* and 26 °C for zebrafish gut isolates, and the matings were carried out on TSA.

Cells from 1 ml of each culture were collected by centrifugation at 8,000 × g for 2 minutes and resuspended in 100 μl PBS. The donor and recipient cultures were mixed in equal parts, spotted onto agar plates and incubated for 16 hours at 26 °C. The media did not contain any antibiotics, nor DAP to prevent *E. coli* AT1036 from proliferating. An equal volume of donor and recipient was also spotted onto separate agar plates. The cells were scraped from the plate using a sterile inoculation loop, resuspended in 1.0 ml PBS buffer, and a dilution series was spread onto differentially selective agar media to enumerate the donors (50 μg.mL^−1^ Km, 10 μg.mL^−1^ Tc, 100 μg.mL^−1^ DAP), recipients (200 μg.mL^−1^ Rif or Nal) and transconjugants (200 μg.mL^−1^ Rif or Nal, 50 μg.mL^-1^ Km, 10 μg.mL^−1^ Tc). Colonies were counted after 2-3 days of incubation at 26°C. Green-fluorescent transconjugants were also verified by excitation at 488 nm and donor, recipient and transconjugant identities were determined by comparing their 16S rRNA gene sequences to known sequences within the RDP database, as described above.

### 4.6 Plasmid persistence assays

The ability of pB10::*gfp* to persist in a host was determined by monitoring the fraction of plasmid-containing cells in a population over 10 days, as described previously (Loftie-Eaton *et al.*, 2016). Briefly, precultures were grown overnight in the presence of kanamycin and tetracycline, and on each subsequent day 4.9 μl of culture was transferred into 5 ml of fresh medium without antibiotics and incubated in a shaking incubator for 24 hours, yielding approximately 10 generations per day. Cultures were spread daily onto nonselective TSA agar such that approximately 100 to 400 colonies were obtained per sample. The fraction of plasmid-containing colonies within each sampled population was determined by counting the fluorescent and non-fluorescent colonies during exposure to a 488-nm light source.

### 4.7 In vivo plasmid transfer

Thirty-two zebrafish of mixed sex were randomly divided into eight groups of four. Each group was put into one of eight tanks (width 10 cm, height 15 cm, depth 25 cm) containing 2 l of filtered water. The eight tanks were divided into two treatment groups in which the fish were fed dry food treated with *E. coli* AT1036 that either contained plasmid pB10::*gfp* (treated tanks, A-D) or was plasmid-free (untreated tanks, E-H). The water was replaced daily with fresh water containing 20 μg.ml^−1^ tetracycline and the fish were fed twice daily with ~35 mg food per tank. The food was prepared every 7 days by suspending the *E. coli* AT1036 with or without plasmid in soybean oil and mixing it with the soy protein-based pellet food (Tetramin) such that the final concentration was ~1 × 10^6^ CFU.mg^−1^ dry food (the CFU count dropped slightly to ~5 × 10^5^ CFU.mg^−1^ during storage over a 9-day period). Thus each 2-l tank received approximately 3 × 10^7^ CFU daily. The fish were fed with the *E. coli*-containing food for 20 days. During the last two days, they were fed with untreated food to minimize *E. coli* donor cells in the gut at the time of harvest. The presence of the donor at that time could confound the transconjugant counts as it may result in plasmid transfer on agar plates rather than in the zebrafish gut.

On the 23^rd^ day, all fish were euthanized and their gut material was harvested aseptically by dissection as described in section 4.3. The gut content from four fish per tank was then pooled and suspended in PBS using the pestle and mortar technique, and bacterial counts were determined by plating a 10-fold dilution series onto different selective TS agar media as follows. Total culturable colony forming units (CFU) were quantified on TSA lacking all antibiotics except cycloheximide (100 μg.ml^−1^). Donor bacteria were quantified based on fluorescence on TSA supplemented with the pB10::*gfp*-selective antibiotics tetracycline (10 μg.ml^−1^), kanamycin (50 μg.ml^−1^), streptomycin (50 μg.ml^−1^) as well as with 100 μM DAP to support the growth of the auxotrophic *E. coli* AT1036 host. Transconjugant bacteria were enumerated on TSA plates supplemented with the same plasmid-selective antibiotics but no DAP (thus counter-selecting the donor). The transconjugant bacteria were distinguishable from the intrinsically resistant gut microbiota based on the fluorescent phenotype encoded by pB10::*gfp*. It should be noted that such transconjugant enumerations will always be an underestimation not only due to limited culturability of all gut bacteria but also because not all bacteria can properly express and fold the fluorescent protein (Cormack *et al.*, 1996).

### 4.8 Zebrafish gut microbiome diversity

After sampling the bacterial cell suspensions from the *in vivo* plasmid transfer experiment for enumeration by plate counting, the remainder of the bacterial cells were collected by centrifugation at 8,000 × g and stored at −20 °C. To account for the bacterial diversity in the water and food, a water control was constructed as follows. Approximately 20 mg of food was mixed with 20 μl of soybean oil and suspended in 2 ml of the system water.

The gDNA was extracted from the zebrafish gut microbiome and water control samples using the two-step enzymatic and bead-beating lysis method described by Yuan *et al.* (2012). Briefly, the frozen cells were thawed on ice and suspended in 900 μl Tris-EDTA buffer (TE; pH 8.0). 50 μl lysozyme (10 mg/ml, Sigma-Aldrich), 6 μl mutanolysin (25 KU/ml, Sigma-Aldrich), and 3 μl lysostaphin (4000 U/ml, Sigma-Aldrich) were added to 250 μl cell suspension and incubated for 1 hour at 37°C. Thereafter, 600 mg of 0.1-mm-diameter zirconia/silica beads (BioSpec) were added to the lysate and the microbial cells were mechanically disrupted using Mini-BeadBeater-96 (BioSpec) at 2100 rpm for 1 minute. The gDNA was purified from the lysate using a QIAamp DNA mini kit (Qiagen). Sequence libraries of the V1 and V3 region of the 16S rRNA genes from each of the samples were constructed in accordance with the Dual Barcoded Two-Step PCR procedure from Illumina (Illumina, 2013). Briefly, using the universal 16S rRNA primers 27F and 534R, the V1-V3 region of 16S rRNA genes was amplified from 2 ng of DNA in 50 μl reactions containing the following components: 1x Standard *Taq* Reaction Buffer (NEB), 3 mM MgCl_2_ (NEB), 0.24 mg.ml^−1^ BSA (Fermentas), 200 μM dNTPs (Fermentas), 50 nM of each of the 27f and 534r primer mixes and 0.025U.μl^−1^ *Taq* DNA polymerase (NEB). The cycling parameters were 95 °C for 2 minutes, 20 cycles of 95 °C for 1 minute, 51 °C for 1 minute and 68 °C for 1 minute, followed by 68 °C 10 minutes and a hold step at 10° C. The PCR products were purified using a QIAquick PCR Purification Kit and visualized on a 1% agarose gel. To attach the barcodes, the PCR products were diluted 15-fold in PCR grade water and 1 μl of each was transferred into 20 μl PCR reaction mix containing 1x Standard *Taq* reaction Buffer, 4.5 mM MgCl_2_, 0.24 mg.ml^−1^ BSA, 75 nM of the barcoded primer, 200 μM dNTPs and 0.05 U.μl-1 *Taq* DNA polymerase. The cycling parameters were 95 °C for 1 minute, 10 cycles of 95 °C for 30 seconds, 60 °C for 30 seconds and 68 °C for 1 minute, followed by 68 °C 5 minutes and a hold step at 10°C. The PCR products were once again purified using a QIAquick PCR Purification Kit and visualized on a 1% agarose gel. The purified, barcoded amplicon libraries were quantified, pooled equimolar and prepared for sequencing on a MiSeq DNA Sequencer (Illumina) by the Genomics Resources Core Facility at the Institute for Bioinformatics and Evolutionary Studies (Moscow, ID) according to their standard operating procedures.

Raw unclipped DNA sequence reads from the Illumina platform were cleaned, assigned and filtered by the Genomics Resources Core Facility using custom scripts, after which the sequence reads were assigned to bacterial taxa using the Naïve Bayesian Classifier for Rapid Assignment of rRNA Sequences (Wang *et al.* 2007) at the Ribosomal Database Project (https://rdp.cme.msu.edu/). The OTU table was interrogated and visualized using R 3.3.0. The 16S RNA gene sequences have been deposited in the Sequence Read Archive at NCBI (Accession: PRJNA601447).

## 5. Acknowledgements

This work was funded in part by the National Institute of Allergy and Infectious Diseases grant R01 AI084918 of the National Institutes of Health (NIH). We are grateful to the IBEST Genomics Research Core for help with DNA sequencing, supported by the Institutional Development Award (IDeA) from the National Institute of General Medical Sciences (NIGMS) of the NIH under grant number P30 GM103324. We also thank Matt Singer for dissecting the zebrafish from the *in vivo* experiment.

